# Genome-scale characterization of wild yeasts reveals cryptic diversity and population structure across three genera

**DOI:** 10.64898/2026.08.06.743242

**Authors:** Kennadi A. Shumaker, Kaitlyn Taylor, Spencer J. Gray, Matthew L. Bochman

**Author notes:** Author for correspondence: Matthew L. Bochman, Molecular & Cellular Biochemistry Department, Indiana University, Bloomington, IN, USA. These authors contributed equally to this work (Kennadi A. Shumaker, Kaitlyn Taylor, Spencer J. Gray).

## Abstract

Environmental surveys of wild yeasts typically rely on ribosomal barcodes, which cannot resolve cryptic species, interspecific gene flow, mixed cultures, or population structure. To determine what genome-scale characterization adds, we sequenced a representative panel of wild yeasts spanning the genera *Saccharomyces*, *Schizosaccharomyces*, and *Lachancea* using Oxford Nanopore long-read whole-genome sequencing and placed each isolate within published reference datasets. Whole-genome analyses revealed biologically important features that barcoding alone could not detect. A shagbark-hickory isolate resolved as a genuine two-species co-culture. An oak-bark isolate proved to be *Schizosaccharomyces versatilis*, a recently reinstated species represented by very few known strains, and its analysis demonstrated that standard assembly-quality benchmarks can be misleading for deep-branching taxa. Three *Lachancea thermotolerans* isolates formed a distinct, previously unsampled population within the wild tree-associated lineage, extending its known geographic range. In contrast, an apparent signal of *Saccharomyces eubayanus* introgression in two beer-associated *S. cerevisiae* isolates disappeared after analysis with matched negative controls and *de novo* assemblies, showing that it reflected mapping artifacts rather than genuine ancestry. Together, these results demonstrate that inexpensive long-read whole-genome sequencing transforms wild-yeast bioprospecting from species identification into a genome-scale framework for resolving cryptic diversity, population structure, and mixed cultures while providing stronger support – and stronger limits – for evolutionary inference.

**SIGNIFICANCE:** Most surveys of wild yeasts identify isolates using short DNA barcodes, which are well suited for naming species but often miss the evolutionary relationships and hidden diversity within them. By applying inexpensive whole-genome sequencing to a diverse collection of environmental yeasts, we uncovered previously undetected mixed cultures, a rare recently recognized species, and a distinct wild population, while also showing that an apparent case of interspecies gene exchange was instead a technical artifact. These results demonstrate that genome-scale analysis can both reveal biological diversity that simpler methods overlook and provide the evidence needed to avoid misleading evolutionary conclusions, making it a powerful new approach for studying natural microbial populations.

## INTRODUCTION

The Declaration of Fermentation (DoF) project at Indiana University Bloomington isolated a single wild *Saccharomyces cerevisiae* strain from the bark of a campus landmark tree, established its wild provenance by whole-genome sequencing and phylogenomic placement, and partnered with local breweries to deploy it in a community-brewed ale (Gray, et al. 2026). Beyond its educational and community aims, DoF produced two durable research assets: a growing collection of wild yeasts sampled from local trees, fruit, and fermentations, and a reproducible approach for situating an environmental isolate genomically among previously described strains. Here, we extend both from a single strain to a representative, genome-sequenced panel.

Wild-yeast collections span the breadth of the yeast subphyla, not merely *S. cerevisiae* (Hittinger 2013; Kurtzman, et al. 2011). The same substrates that yield wild *Saccharomyces* also yield other budding yeasts such as *Lachancea* and the early-diverging fission yeasts (*Schizosaccharomyces*) (Osburn, et al. 2018), whose deep phylogenetic position within the Taphrinomycotina makes them both evolutionarily distinctive and poorly represented in reference resources (Kurtzman and Sugiyama 2015; Liu, et al. 2009; Sipiczki 1995). Each of these lineages poses questions that ribosomal barcode markers (the ITS region and the D1/D2 domain of the large-subunit rRNA gene (Kurtzman and Robnett 1997; Schoch, et al. 2012)) cannot answer: barcodes do not resolve species within the *Saccharomyces sensu stricto* complex (Hittinger 2013; Libkind, et al. 2011), do not detect interspecific hybridity or introgression (Borneman, et al. 2016; Hittinger 2013), and readily miss cryptic, recently delimited, or co-cultured taxa (Bickford, et al. 2007; Lucking, et al. 2021). Long-read whole-genome sequencing, now inexpensive enough to apply isolate by isolate (Warburton and Sebra 2023), replaces these blind spots with genome-scale placement against reference strains *via* average nucleotide identity (ANI) for species assignment (Shaw and Yu 2023), whole-genome phylogenomics for population placement, and direct comparison to type-strain assemblies for structural and introgression analysis (Dujon 2010; Hittinger 2013; Warburton and Sebra 2023).

Three cases from our own collection (Osburn, et al. 2018) make this concrete. One isolate, YH156, whose ribosomal barcode suggested that it was *Schizosaccharomyces japonicus*, was ultimately found to be *Schizosaccharomyces versatilis*, a species very recently reinstated from *S. japonicus* var. *versatilis* (Brysch-Herzberg, et al. 2024). Our isolate is among the very few reported for this lineage and adds to newly recognized within-species structure, while its characterization exposes repeat- and reference-database pitfalls that would mislead any pipeline calibrated on model yeasts. In *Saccharomyces*, wild *S. cerevisiae* isolates distribute across multiple lineages of the global population (Loegler, et al. 2024), and the two isolates recovered from spontaneously fermented beer illustrate the opposite lesson: an apparent interspecific introgression signal that proved, under matched negative controls, to be indistinguishable from the false-positive floor of the detection method itself (companion paper, Taylor *et al*.). In *Lachancea*, we place our *L. thermotolerans* isolates relative to the described diversity of this increasingly studied species (Vicente, et al. 2025).

Together these three cases make a single point: extending place-based wild-yeast bioprospecting from one strain (Gray, et al. 2026) to a genomically characterized, taxonomically broad panel – and placing each isolate against the strains already known – turns a catalog of names into a view of cryptic species, interspecific gene flow, and population structure across the yeast tree.

## RESULTS

### A three-genus wild-yeast panel placed against described strains

As a follow-up to the DoF project (Gray, et al. 2026), we assembled a panel of nine wild-yeast isolates recovered from tree bark, fruit, and spontaneously fermented (“wild”) beer in the United States, together with the campus burr oak isolate *S. cerevisiae* DoF1 (Table 1). Each isolate was sequenced by Oxford Nanopore long-read whole-genome sequencing and characterized with a single uniform pipeline (Figure 1): ribosomal barcode extraction (ITS and the D1/D2 domain of the LSU rRNA gene), whole-genome ANI against type and reference strains, and – wherever the barcode could not resolve the assignment – genus-specific phylogenomic placement against a published population panel.

**Figure 1.**
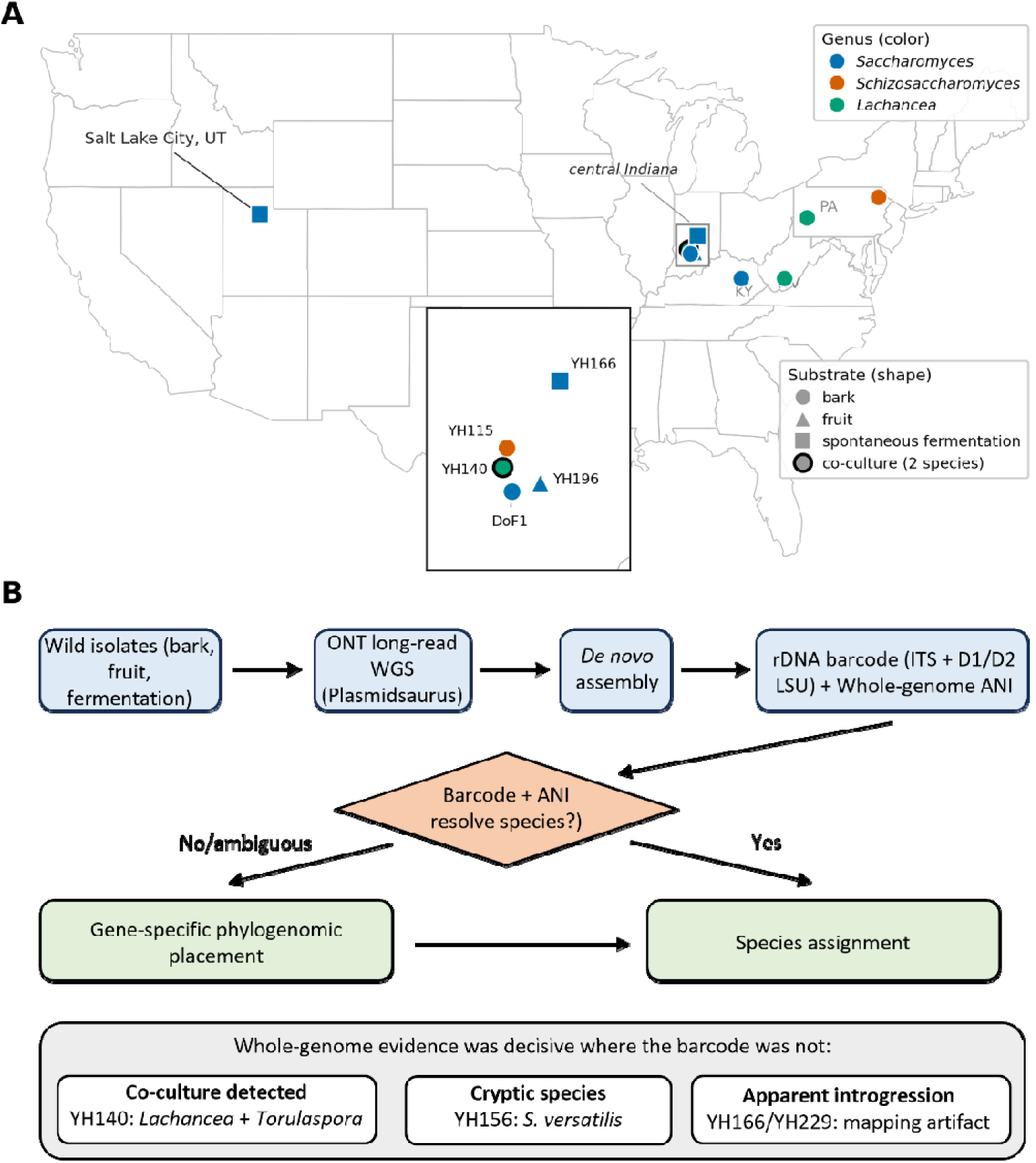
A wild-yeast bioprospecting panel and the identification-placement pipeline. (**A**) Geographic origin of the panel isolates across the United States, with markers colored by genus (*Saccharomyces*, *Schizosaccharomyces*, *Lachancea*) and shaped by substrate (bark, fruit, spontaneous fermentation). (**B**) The uniform per-isolate workflow: Oxford Nanopore long-read whole-genome sequencing → *de novo* assembly → ribosomal barcode extraction (ITS + D1/D2 LSU) → whole-genome ANI against type/reference strains → genus-specific phylogenomic placement for isolates the barcode could not resolve. The bottom highlights the three cases where whole-genome evidence overturned or refined the barcode call: the co-culture (YH140), the cryptic species (YH156), and the beer isolates, whose apparent *S. eubayanus* introgression proved on controlled analysis to be a mapping artifact (YH166/YH229).

**Table 1.** Isolate collection metadata and final species assignments.

| Isolate | Substrate | Locality | Collection date | Collector | Final identification |
| --- | --- | --- | --- | --- | --- |
| YH26 | Tulip poplar bark | New River Gorge, WV | 17 May 2014 | K. Brown | *Lachancea thermotolerans* |
| YH72 | Ash bark | New Kensington, PA | 8 Jul 2014 | M. L. Bochman | *Lachancea thermotolerans* |
| YH115 | Oak bark | Bloomington, IN | 23 Jan 2015 | K. Osburn | *Schizosaccharomyces pombe* |
| YH123 | Oak bark | Red River Gorge, KY | 2 May 2015 | M. L. Bochman | *Saccharomyces cerevisiae* |
| YH140 | Shagbark hickory bark | Bloomington, IN | Jun 2015 | K. Osburn | *L. thermotolerans* + *Torulaspora delbrueckii* |
| YH156 | Oak bark | I-81 rest area, eastern PA | 11 Jun 2015 | M. L. Bochman | *Schizosaccharomyces versatilis* (nom. inval.) |
| YH166 | Wild-fermented beer | Indianapolis, IN | Summer 2015 | J. Miller | *Saccharomyces cerevisiae*<sup>a</sup> |
| YH196 | Pawpaw fruit | Bloomington, IN | Sep 2015 | D. Nickens | *Saccharomyces cerevisiae* |
| YH229 | Wild-fermented beer | Salt Lake City, UT | 2015 | Anonymous homebrewer | *Saccharomyces cerevisiae*<sup>a</sup> |
| DoF1 | Burr oak bark | Bloomington, IN | Feb 2026 | Gray *et al.* | *Saccharomyces cerevisiae*<sup>b</sup> |

<sup>a</sup> YH166 and YH229 were initially flagged as putative *S. cerevisiae* × *S. eubayanus* hybrids on the basis of reduced read-mapping to the S288C reference, and subsequently as carrying minor *S. eubayanus* nuclear introgression. Neither call survived testing against depth-matched negative controls (see *Screening for interspecific S. eubayanus ancestry*): the apparent interspecific fraction fell at or below the false-positive floor of the mapping method, and no *S. eubayanus* sequence is present in either de novo assembly. Both are reported here as *S. cerevisiae* without detectable introgression; the artifact is characterized in the companion study [cite: companion methods paper].
<sup>b</sup> DoF1 was isolated and its wild provenance established in the Declaration of Fermentation study (Gray *et al.* 2026); full details are reported there. It is included here in the population-genomic placement analysis.

This resolved five species distributed across three genera: *Saccharomyces* (five *S. cerevisiae* = YH123, YH166, YH196, YH229, and DoF1), *Schizosaccharomyces* (YH115, *S. pombe*; and YH156, *S. versatilis*), and *Lachancea* (YH26 and YH72, *L. thermotolerans*), with *Torulaspora delbrueckii* additionally recovered from one *Lachancea* co-culture (YH140; Table 1). In three cases the ribosomal barcode alone was insufficient, and whole-genome evidence was decisive. Isolate YH140 proved not to be a single strain but a two-species co-culture: contig binning cleanly partitioned its assembly into a *L. thermotolerans* component (53.3% of the assembly) and a *T. delbrueckii* component (46.2%), each recovered as a coherent genome bin (Supplementary Figure S1). The two beer-associated *S. cerevisiae* strains YH166 and YH229 carried an interspecific signal invisible to barcoding, and the fission-yeast isolate YH156 was initially mis-assignable at the barcode level (Table 1). We treat each genus in turn below, in every case placing our isolates relative to the strains already described.

### *Saccharomyces*: diverse wild *S. cerevisiae* and an interspecific signal that did not survive

Placed within a global panel of 3,034 *S. cerevisiae* genomes by maximum-likelihood inference from 46,705 genome-wide SNPs (Loegler, et al. 2024), the five *S. cerevisiae* strains in the collection distributed across multiple ecological lineages rather than forming a single group (Figure 2). YH123, isolated from oak bark in the Red River Gorge (KY), and the campus burr oak (*Quercus macrocarpa*) isolate DoF1 placed together within the Wild clade, consistent with their wild tree bark provenance (Table 1) and with the wild status established for DoF1 in the source study (Gray, et al. 2026). Of the two beer isolates, YH229 placed within the Beer superclade – toward its edge in the coarse placement – but resolves at finer scale to a single, well-supported sub-clade (*12. Belgium Beer 1*). Its nearest neighbors agree unanimously at both the immediate-sister radius (3/3) and the grandparent radius (4/4), each at maximum FastTree support (1.0). YH166, by contrast, did not resolve cleanly at any scale (see below). The pawpaw-fruit (*Asimina triloba*) isolate YH196 likewise did not resolve into a named lineage, falling among the panel’s Unassigned/Admixed isolates. Its single nearest neighbor belongs to a French Guiana-associated lineage, but nearest-neighbor identity within this diffuse, low-support group is not itself informative of shared ancestry, as the same pattern is observed for YH166 (see below).

**Figure 2.**
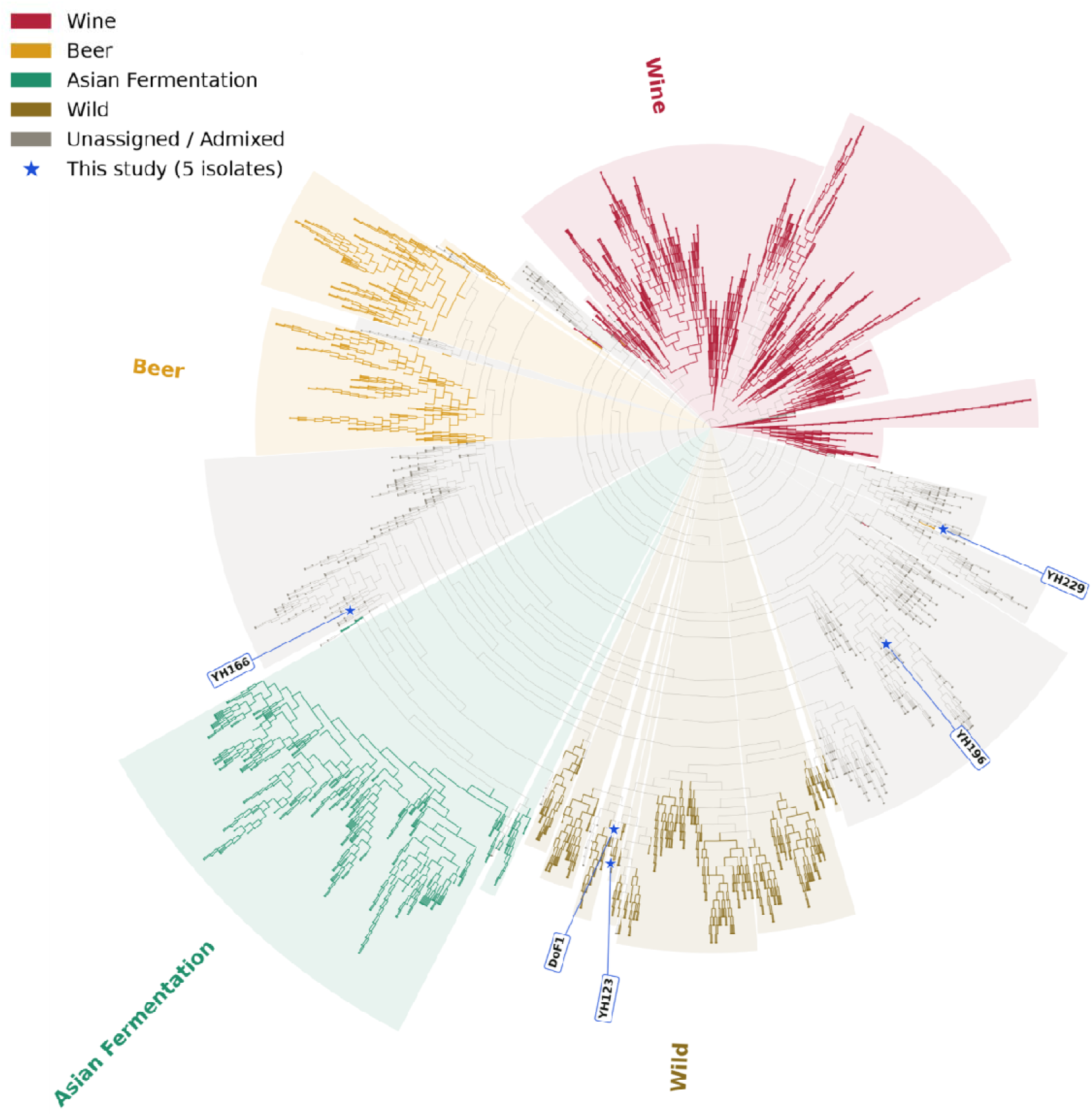
Phylogenomic placement of the collection’s *S. cerevisiae* strains within global diversity. Circular maximum-likelihood cladogram (FastTree) of *S. cerevisiae* YH123, YH166, YH196, YH229, and DoF1 placed among the 3,034-genome global panel (Loegler, et al. 2024), inferred from 46,705 genome-wide SNPs shared across all samples. Panel tips are shaded by ecological category (Wine, Beer, Asian Fermentation, Wild, Unassigned/Admixed); the five study isolates are marked with stars and labeled. YH123 and DoF1 place together in the Wild clade. YH229 places at the periphery of the Beer clade and YH166 at the edge of the Asian-fermentation clade. The pawpaw isolate YH196 falls in an unassigned/admixed position outside the named lineages.

YH166’s *S. cerevisiae* sub-genome did not resolve into any panel clade at the coarse level (Figure 2), and (unlike YH229) this ambiguity persists at finer scale. Its nearest neighbors at both the immediate-sister and grandparent radii are themselves individually unassigned/admixed isolates from unrelated geography (China, Italy, Finland), at correspondingly low support (0.838 and 0.846). A well-supported grouping emerges only at a much broader radius (383 tips, FastTree support 0.996), of which 305 (80%) classify as Asian Fermentation. Consistent with this, a superclade-level allele-sharing analysis at YH166’s 35,800 heterozygous sites shared with the panel catalog found the highest mean alternate-allele frequency in the Asian Fermentation superclade (0.4625), followed by Wild (0.3606), Beer (0.3497), Unassigned/Admixed (0.3247), and Wine (0.2935); no superclade approached fixation. Together, these indicate genuine admixture with an Asian Fermentation lean, not a resolved single-lineage origin. This characterization rests entirely on SNP-based population placement and is independent of the *S. eubayanus* interspecific screen described below.

The two beer-associated isolates, YH166 and YH229, initially produced a strong signal suggestive of *S. cerevisiae* × *S. eubayanus* hybridity: reduced and uneven read mapping against the *S. cerevisiae* S288C nuclear reference (60.5% and 77.6% primary mapping, respectively, *vs*. 93-96% for the other *S. cerevisiae* isolates). That signal did not survive controlled analysis. We report its collapse in full, because the sequence of corrections is more informative than the negative result alone.

Supplying the reference with the *S. cerevisiae* mitochondrion and 2-micron plasmid – both absent from the standard S288C nuclear assembly, leaving reads from those high-copy elements with no legitimate target – removed most of the apparent interspecific signal and reduced the picture from balanced hybridity to a minor, patchy nuclear introgression (companion paper, Taylor *et al*.). Testing that residual against matched negative controls removed the remainder. Two wild *S. cerevisiae* from the same collection and sequencing platform with no plausible *S. eubayanus* contact (YH123 and DoF1) were processed through the identical pipeline. At sequencing depth matched between isolate and control, the apparent *S. eubayanus* fraction in YH166 and YH229 fell at or below the fraction recovered from those controls. Most candidate blocks recurred at the same coordinates in the controls. The interior, non-subtelomeric signal in YH229 fell to zero at every matched depth, while the controls retained more. The sub-diploid relative dosage we had taken as supporting evidence was reproduced by the controls as well. Per-read realignment showed that the most depth-robust candidate blocks comprised either reads whose primary alignments lay on *S. cerevisiae* chromosomes or reads aligning at higher identity to *S. cerevisiae* than to *S. eubayanus*. Finally, no contig of either *de novo* assembly mapped to *S. eubayanus* anywhere in the genome, though both assemblies are near the expected *S. cerevisiae* genome size and contiguity (YH166, 12.40 Mb in 41 contigs, N50 801 kb; YH229, 12.25 Mb in 31 contigs, N50 822 kb).

We therefore report YH166 and YH229 as wild beer-associated *S. cerevisiae* with no detectable *S. eubayanus* introgression. Both carry *S. cerevisiae*-type mitochondria and 2-micron plasmids, and neither shows any *S. cerevisiae*/*S. eubayanus* chimeric junction – a whole-assembly scan recovered no single contig joining *S. cerevisiae* and *S. eubayanus* nuclear sequence (0 of 25 contigs in YH229, 0 of 39 in YH166) – consistent with the assembly result above. Their population-level placements, described above, are unaffected by this result.

### *Schizosaccharomyces*: a rare isolate of the recently reinstated species *S. versatilis*

Isolate YH156, previously reported to be *S. japonicus* (Osburn, et al. 2018; Peepall, et al. 2019), initially resembled a failed sequencing run. While its raw read quality and yield were comparable to the other eight isolates, its assembly collapsed to 426 kb in 54 contigs and scored 0% BUSCO completeness against the budding-yeast (Saccharomycetes) ortholog set applied to the rest of the batch. Two compounding causes, not a sequencing failure, were responsible. First, the 0% score reflected the wrong benchmark: BLAST of the assembled contigs placed YH156 in the Taphrinomycotina, an early-diverging subphylum against which a budding-yeast gene set cannot score (Manni, et al. 2021; Shen, et al. 2018). Second, a ribosomal RNA repeat array at ∼1,500× coverage inflated the genome-wide mean so far that correctly assembled single-copy sequence at ∼300× was discarded by the assembler’s low-coverage filter. Approximately 2.5 Mb of legitimate sequence was recoverable from the pre-filter output. Barcode and ANI analysis then reassigned the isolate decisively: YH156 shares 99.14% ANI with the *Schizosaccharomyces versatilis* type strain CBS 103 (assembly GCA_032882995.1) but only 87.94% with *S. japonicus* (GCF_000149845.2), consistent with its assignment to *S. versatilis* following the recent reinstatement of that species (Brysch-Herzberg, et al. 2024).

This assignment is notable because YH156 is now one of very few isolates of *S. versatilis*, which is currently only represented by the type strain CBS 103 (Wickerham and Duprat 1945) and the related genetic strain CBS 5679 (Etherington, et al. 2024). Thus, YH156, a wild environmental isolate from tree bark, is among the first such isolates reported for this lineage and adds a new data point to the within-species structure that has only recently been recognized.

Regardless, reassembly confirmed that the fragmentation was an artifact. Across three assemblers, Flye failed to resolve chromosome 2 under any configuration tested (1.2% reference coverage, *vs*. 91.8% for Canu and 95.1% for hifiasm) (Figure S2). Canu and hifiasm each independently produced a contig fusing two chromosome ends at identical coordinates, *i.e.*, two algorithms converging on the same junction, confirming a real, repeat-driven misassembly rather than random error. The Canu assembly (highest contiguity, lowest duplication) was split at the verified breakpoint and consensus-polished, yielding a 17,065,046-bp draft in 213 contigs. Against the correct reference (∼15.5 Mb across three chromosomes of 4.46, 7.34, and 3.69 Mb), it recovered near-complete chromosome-scale coverage (93.7, 93.7, and 94.3%). Apparent gene-completeness was itself informative: YH156 recovered 55.4% of the ascomycota_odb12 BUSCO set (Figure S3), a figure that would ordinarily signal a deficient assembly (Manni, et al. 2021). However, the finished same-species reference (*S. versatilis* CBS 103 (Brysch-Herzberg, et al. 2024)) scored essentially identically on the same lineage set (57.9%; with the congener *S. japonicus* at 57.1%), so the orthologs scored “missing” reflect a lineage-database ceiling for these early-diverging fission yeasts, not incompleteness of the YH156 assembly. This calibration is a concrete caution against applying model-yeast completeness expectations to deep-branching taxa.

Whole-genome alignment of YH156 to the CBS 103 type-strain chromosomes (CP130697.1–CP130699.1), performed without reference-guided scaffolding so as not to mask real differences, recovered broadly conserved synteny alongside localized rearrangements, and identified the Tf1/Tf2 LTR retrotransposon families characteristic of this lineage (Figure 3). Rearrangement calls concentrated away from contig boundaries, arguing for real signal rather than fragmentation noise. We report three high-confidence events (Figure S4): an ∼768 kb inversion on chromosome 1 (∼2.16–2.93 Mb), two adjacent inversions on chromosome 2 (∼295 and ∼115 kb, ∼6.02–6.46 Mb), and an intra-chromosomal translocation of ∼294–357 kb on chromosome 3 (∼3.17–3.53 Mb). A dense duplication/translocation signal on chromosome 2 (∼2.4–5.5 Mb) is likely dominated by repetitive sequence and is deliberately excluded pending higher-contiguity data. Given the draft, contig-level nature of the assembly, we treat individual structural calls as provisional. The panel’s second fission-yeast isolate, YH115 (oak bark, Bloomington IN), was a straightforward *S. pombe* by barcode and ANI, and is reported without further structural analysis.

**Figure 3.**
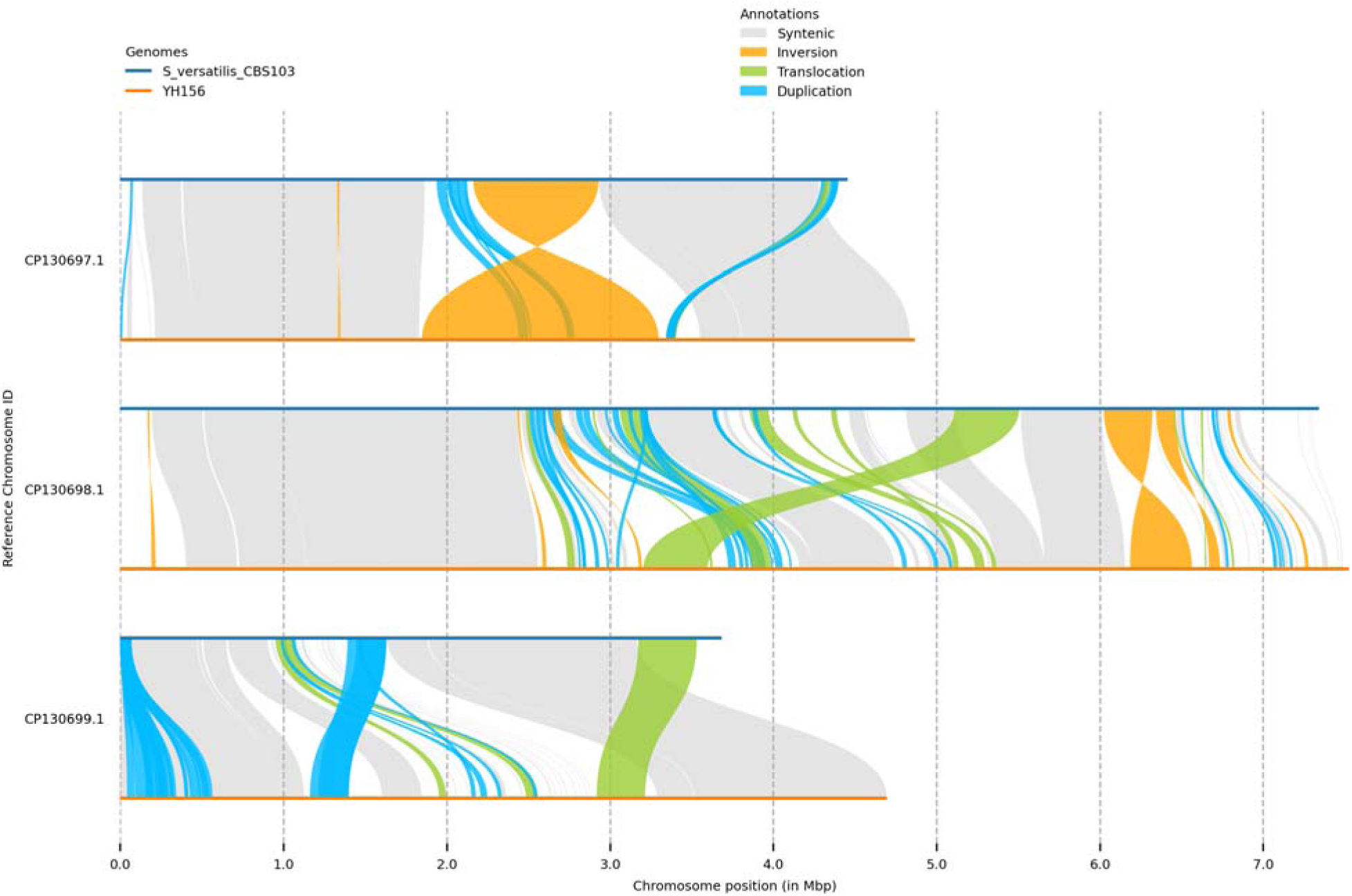
Whole-genome synteny of *Schizosaccharomyces versatilis* isolate YH156 to the type strain CBS 103. SyRI/plotsr comparison of the YH156 draft assembly (orange) against the *S. versatilis* CBS 103 type-strain chromosomes (blue; CP130697.1, CP130698.1, CP130699.1), performed without reference-guided scaffolding. Syntenic regions (grey) and rearrangements are color-coded per SyRI: inversions (orange), translocations (green), and duplications (blue). The three high-confidence events discussed in the text are the ∼768 kb chromosome-1 inversion (∼2.16–2.93 Mb), the two adjacent chromosome-2 inversions (∼295 and ∼115 kb, ∼6.02–6.46 Mb), and the ∼294–357 kb intra-chromosomal chromosome-3 translocation (∼3.17–3.53 Mb). The dense chromosome-2 signal near ∼2.4–5.5 Mb is likely repeat-driven and is excluded pending higher-contiguity data. Because the YH156 assembly is a contig-level draft, individual rearrangement calls are provisional, and rearrangement calls concentrate away from contig boundaries.

### *Lachancea*: a distinct wild sub-population within the tree-associated lineage

The three tree bark-associated *L. thermotolerans* representatives of the collection – YH26 (tulip poplar (*Liriodendron tulipifera*), WV), YH72 (ash (*Fraxinus excelsior*) PA), and the *L. thermotolerans* bin recovered from the YH140 co-culture (shagbark hickory (*Carya ovata*), IN; 57,944 reads) – were placed against the 145-strain *L. thermotolerans* population panel of (Vicente, et al. 2025), which resolves six ecological/geographic clusters. All three isolates assigned unambiguously and concordantly to the Canada-trees cluster, the panel’s wild, tree-associated North American lineage (Figure S5). Supervised ADMIXTURE (K = 6) estimated 99.995% Canada-trees ancestry for the isolates. Principal-component analysis (PC1, 21.80%; PC2, 19.40% of variance) placed them nearest the Canada-trees strains, and a genome-wide BIONJ tree recovered all three among Canada-trees members (Figure 4). For wild bark isolates, membership in the tree-associated cluster is the ecologically expected outcome, and the concordant placement of the YH140 bin alongside the two clonal isolates confirms that binning recovered a genuine *L. thermotolerans* genome.

**Figure 4.**
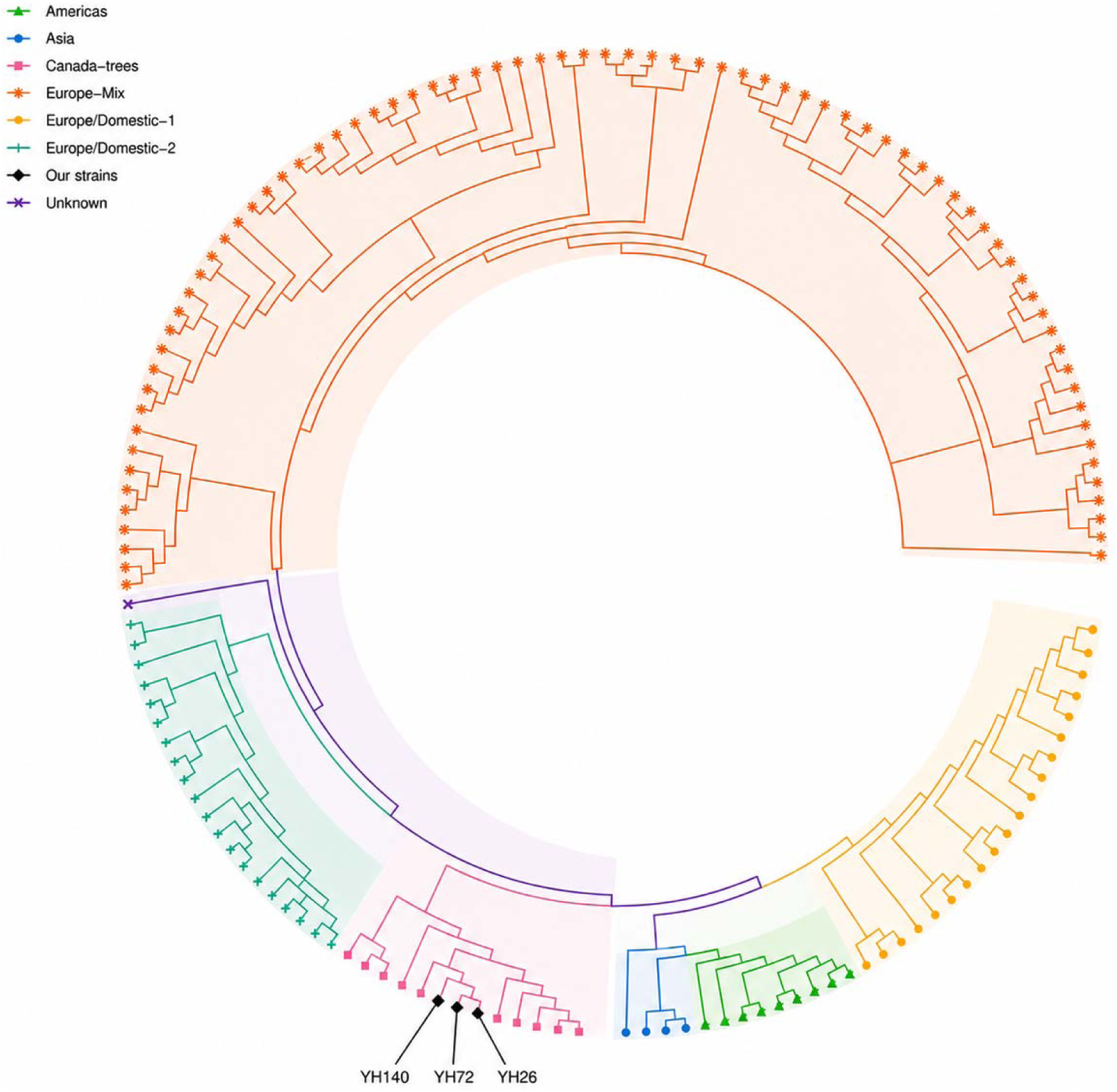
BIONJ tree of the reconstructed *L. thermotolerans* panel. Genome-wide-distance BIONJ tree of the 145 panel strains (Vicente, et al. 2025) plus the three study strains (ape v5.8-1). The three study strains group among Canada-trees members. Note that, in this reconstruction, the Canada-trees cluster alone was not strictly monophyletic; the tree is therefore presented as corroborating the ADMIXTURE and PCA placement (Figure S5) rather than as an independent topological claim.

The isolates do not, however, merely fall inside the described diversity; they extend it. The three were roughly an order of magnitude closer to one another (pairwise IBS-based distances 0.023-0.032) than to their nearest panel neighbors (0.128-0.134, mean 0.131) – in each case, a Canada-trees strain and the same one (ERR2886527) for two of the three isolates – computed *via* SNPRelate’s individual dissimilarity analysis on 1,015,261 genome-wide SNPs across the 148-sample panel, and were offset from the sampled Canada-trees strains along PC2 (Figure S5). Thus, they constitute a tight, previously unsampled wild population within the Canada-trees lineage, extending its documented range into the eastern and midwestern United States (the existing cluster spans Canadian and southern-US wild strains) (Vicente, et al. 2025). This is the *Lachancea* arm’s contribution to the study’s through-line: not a surprising placement, but genuinely new diversity nested within a known lineage.

Two caveats bound these conclusions, though the first has since been substantially strengthened. First, because the source study’s filtered genotype matrix (Vicente, et al. 2025) was not available when this analysis was initially performed, the reference panel was reconstructed from published raw reads. The reconstruction retained 93.0% of SNPs *vs*. the 93.9% reported by the source study and used bcftools-native filter statistics as analogs of the original GATK-based ones. Thus, it was methodologically comparable but not a byte-for-byte reproduction of the published calls. The source study’s authoritative filtered VCF was subsequently obtained and used to independently re-run the ADMIXTURE and PCA placement analyses against the real published calls: both were concordant with the reconstruction (99.995% Canada-trees ancestry in both; PCA placement nearest the Canada-trees centroid in both, with nearly identical variance explained). Figure 4 reports the reconstruction-based results. Second, in the reconstructed tree, the Canada-trees cluster (uniquely among the six) was not strictly monophyletic. The same non-monophyly recurred when the tree was independently rebuilt against the published filtered VCF, indicating this reflects genuine structure within the cluster rather than a reconstruction artifact. The tree is therefore best read as corroborating the ADMIXTURE and PCA placement rather than as an independent topological claim. The placement itself is robust across all three methods, and now across both the reconstructed and published reference panels.

## DISCUSSION

A yeast collection built from ordinary substrates, largely sampled near a single campus, was not designed to test any one hypothesis, and did not need to. Examined at whole-genome resolution rather than by barcode alone, each of its three genera returned the same lesson. In *Saccharomyces*, whole-genome analysis overturned a conclusion that read mapping alone had supported; in *Schizosaccharomyces*, a cryptic species reported from only a handful of prior isolates; in *Lachancea*, a wild population differentiated from anything yet sequenced. Read together, they argue that inexpensive long-read whole-genome sequencing turns environmental yeast bioprospecting from a cataloguing exercise into an instrument for detecting cryptic speciation, mixed culture, and population structure across the yeast tree – and that the same data are what discipline the inferences drawn from them.

### What genome-scale placement recovers that barcodes cannot

The clearest through-line of this study is methodological. Ribosomal markers assign genus reliably and, for most fungi, species (Kurtzman and Robnett 1997; Schoch, et al. 2012).

However, they are blind to precisely the features that carry evolutionary information in this clade. A single co-culture (YH140) and a species-level misassignment for a deep-branching fission yeast (YH156) went unnoticed and were misreported on barcode evidence alone (Osburn, et al. 2018). Here, both were resolved only by whole-genome ANI and phylogenomic placement against reference strains. The beer isolates make the complementary point: whole-genome evidence was decisive there too, but in the direction of withdrawing a claim rather than making one. As long-read sequencing of a fungal genome approaches the cost of a handful of Sanger reads, there is diminishing justification for identifying environmental fungal isolates by ribosomal barcode alone.

### A lineage-database ceiling for deep-branching taxa

The characterization of YH156 illustrates a trap that grows more acute the further an isolate sits from model lineages. Its 55.4% BUSCO completeness would, by the conventions applied to *Saccharomyces*, flag a badly deficient assembly (Figure S3). However, the finished *S. versatilis* type strain and *S. japonicus* itself scored barely higher on the same ortholog set, placing the “missing” fraction not in the YH156 assembly but in the mismatch between a benchmark curated on well-sampled taxa and the true gene content of early-diverging fission yeasts.

Completeness scores are routinely used as gatekeeping filters and as evidence of assembly quality. Our calibration shows that for deep-branching taxa, such thresholds can be actively misleading, and that the appropriate control is a finished conspecific or close relative run through the identical benchmark rather than an absolute cutoff. This caution generalizes well beyond *Schizosaccharomyces* to any genomic survey reaching into under-sampled regions of a phylogeny. That YH156 is itself a rare isolate of *S. versatilis* – a species reinstated from *S. japonicus* var. *versatilis* only recently (Brysch-Herzberg, et al. 2024) and represented by extremely few strains – adds a modest but real contribution to the population sampling of a taxon whose within-species structure has only just begun to be described.

### An interspecific signal that did not survive

The two beer-associated *S. cerevisiae* were, at the outset, the study’s most familiar-looking result. Interspecific *S. eubayanus* ancestry in beer-associated *Saccharomyces* is well documented, from the lager hybrids onward (Langdon, et al. 2019; Libkind, et al. 2011), and both isolates presented exactly the mapping signature such ancestry produces. The instructive part is that the signal survived three rounds of analysis and failed the fourth. Competitive mapping against a nuclear-only combined reference indicated balanced hybridity. Supplying the reference with the *S. cerevisiae* mitochondrion and 2-micron plasmid reduced that to minor, patchy introgression. Only a matched negative control – an unrelated wild *S. cerevisiae* from the same collection, sequencing chemistry, and depth regime, run through the identical pipeline – showed that what remained was not distinguishable from what the method recovers in a genome that cannot contain it.

Three features of that failure are worth stating plainly, because each contradicts an assumption we ourselves held. First, the apparent introgression grew with sequencing depth rather than stabilizing, so YH229’s larger signal reflected its unusually deep coverage rather than more introgressed sequence. Second, long reads did not protect against it: the artifact concentrates near chromosome ends, where sequence similarity between these species is genuine rather than a repeat-spanning problem that read length can solve. Third, sub-diploid relative dosage, which we had treated as positive evidence that the introgressed material was real and subclonal, is reproduced by artifactual windows in control strains, because low-uniqueness regions mechanically yield reduced unique-read depth.

We report this at length rather than simply omitting the isolates, because the near-miss is the transferable result. Any pipeline that quantifies interspecific nuclear ancestry by competitive read mapping is exposed to the same failure mode, and the error inflates precisely the signal such screens exist to detect. The practical safeguards are inexpensive: 1) include organellar and plasmid sequence in the reference or mask it explicitly; 2) run a known-negative control at matched sequencing depth through the identical pipeline; 3) test whether candidate blocks recur at shared coordinates in unrelated strains; and 4) confirm that candidate introgressed sequence is present in the *de novo* assembly, not only in the read mapping. The mechanisms, their quantification, and their consequences for published introgression calls are developed in a companion study (companion paper, Taylor *et al*.).

### *Lachancea*: concordant ecology and an under-sampled wild population

The *L. thermotolerans* isolates gave the one result that met, rather than overturned, the naïve expectation and are the more useful for it. Wild bark isolates placing in the panel’s wild, tree-associated (“Canada-trees”) lineage (Vicente, et al. 2025) is the concordance between ecology and genotype one hopes for, and the co-clustering of the metagenomically binned YH140 genome with the two clonal isolates is a satisfying internal check on the binning. That said, the three isolates were an order of magnitude more similar to one another than to any sequenced panel strain and offset from them in ordination space, marking a discrete wild population within that lineage that current panels have not captured and extending its documented range into the eastern and midwestern United States (Figure 4 and S5). Even a well-constructed 145-strain panel, then, leaves conspicuous geographic gaps in the wild diversity of a species otherwise studied intensively for its winemaking relevance (Vicente, et al. 2025), a reminder that anthropized sampling and wild diversity are not the same target.

### Limitations

Several constraints bound these conclusions. The panel is small (nine isolates plus the previously reported DoF1 (Gray, et al. 2026)), so ecological associations are illustrative rather than statistical. The YH156 assembly is draft-quality and contig-level, so its structural-variant and synteny calls are provisional, and its finishing would benefit from long-range scaffolding. The negative introgression finding for YH166 and YH229 establishes that no *S. eubayanus* ancestry is detectable above the method’s false-positive floor. It does not exclude introgressed segments smaller than that floor, and one subtelomeric locus that failed only the positional and assembly criteria is reported in the companion study (companion paper, Taylor *et al*.) as the single locus the data cannot fully exclude. The *Lachancea* placement was built from a panel we reconstructed from raw reads rather than the source authors’ filtered genotypes; the reconstruction is closely concordant with the original pipeline (93.0% *vs*. 93.9% SNP retention). We subsequently obtained the source study’s authoritative filtered VCF and independently re-ran the ADMIXTURE and PCA placement analyses against the real published calls: both were concordant with the reconstruction (99.995% Canada-trees ancestry in both; PCA placement nearest the Canada-trees centroid in both, with nearly identical variance explained). The reconstructed tree’s non-monophyletic recovery of one cluster means we lean on this ADMIXTURE and PCA evidence, now doubly confirmed, for that inference. Phylogenetic support was assessed with approximate (SH-like) rather than model-based bootstrap methods.

### Outlook

This work is a proof of concept for a particular kind of science: locally sampled, community-connected bioprospecting (Gray, et al. 2026) coupled to genome-scale placement against the growing public catalogue of yeast diversity, with the identification and placement pipeline released openly [cite: pipeline Zenodo DOI] so that the approach transfers to other collections. Each genus points to a natural follow-up: 1) a dedicated treatment of the *S. versatilis* isolate with a finished assembly and expanded fission-yeast sampling; 2) a wild-*Saccharomyces* introgression study built on negative controls and a *S. eubayanus* panel adequate to detect genuine low-level gene flow below the floor characterized here; and 3) deeper *L. thermotolerans* sampling across North American wild substrates to place the population reported here in its full geographic context. More broadly, as whole-genome characterization of environmental isolates becomes routine, the boundary between a strain survey and an evolutionary-genomics study continues to erode. The same sequencing that identifies a wild yeast now also situates it, and as the beer isolates show, constrains what may responsibly be claimed about it.

## MATERIALS AND METHODS

All bioinformatic analyses were run within conda/bioconda environments (channels conda-forge and bioconda, channel_priority strict) on the Indiana University Quartz HPC cluster. Software versions are given at first mention and collated below.

### Isolates and sampling

Nine yeast isolates from a wild/environmental bioprospecting collection were characterized (Table 1). Isolates were recovered from tree bark, fruit, and spontaneously (“wild”) fermented beer sampled across the eastern and western United States between 2014 and 2015. An additional wild *Saccharomyces cerevisiae* isolate, DoF1 – isolated from the bark of a campus landmark tree at Indiana University Bloomington and shown to be of wild provenance in the Declaration of Fermentation study ((Gray, et al. 2026); NCBI BioProject PRJNA1477971, SRA run SRR39135318) – was included in the population-genomic placement analysis.

### Sequencing

All nine isolates were sequenced by Plasmidsaurus (Louisville, KY, USA) using Oxford Nanopore long-read whole-genome sequencing. Libraries were prepared amplification-free with Oxford Nanopore v14 (kit-14) chemistry using sequence-independent, tagmentation-based fragmentation and a primer-free protocol, sequenced on R10.4.1 flow cells, and delivered as FASTQ. Reads were basecalled with the super-accuracy model dna_r10.4.1_e8.2_400bps_sup@v4.3.0 (R10.4.1 flow-cell chemistry, Dorado super-accuracy basecalling); the model tag was read directly from the raw read headers for YH123, YH166, YH196, and YH229 and is assumed identical for the remaining isolates.

### Part A - Species identification Read quality control and filtering

Raw read statistics were summarized with seqkit v2.13.0 (stats -a) (Shen, et al. 2016). Read-length and quality-score distributions were visualized with NanoPlot v1.47.1 (De Coster, et al. 2018). Reads were quality- and length-filtered with chopper v0.13.0, retaining reads with mean quality Q ≥ 10 and length ≥ 1000 bp.

#### Genome assembly

Filtered reads were assembled *de novo* with Flye v2.9.6-b1802 (Kolmogorov, et al. 2019) in high-quality Nanopore mode (--nano-hq) with an expected genome-size hint of 12 Mb.

#### Assembly completeness assessment

Assembly completeness was assessed with BUSCO v5.8.3 in genome mode (Manni, et al. 2021), with gene models generated by the automatically selected predictor (miniprot v0.18-r281; (Li 2023)) and scored with hmmsearch v3.4. For isolates in the class Saccharomycetes, the saccharomycetes_odb10 lineage dataset (2024-01-08 build; 76 genomes; 2,137 BUSCO groups) was used. Because saccharomycetes_odb10 is calibrated for the class Saccharomycetes and is not an appropriate benchmark for taxa outside it, a lineage correction was applied to isolates found to lie elsewhere: YH115 (*Schizosaccharomyces pombe*) initially scored an implausibly low 61.3% against saccharomycetes_odb10 and was re-assessed against the broader ascomycota_odb10 lineage (365 genomes; 1,706 BUSCO groups; no dedicated Schizosaccharomycetes or Taphrinomycotina lineage exists in the current BUSCO catalog), returning 80.0%, which is the reported value. The same reasoning was applied to the fission-yeast isolate YH156 (see Part B).

#### Ribosomal marker gene identification and extraction

Ribosomal RNA genes (18S, 5.8S, 28S) were annotated with barrnap v1.10.6 (--kingdom fun; this barrnap build supports only bacterial, archaeal, and fungal kingdom models, and fun is the taxonomically correct choice for all isolates here) (Seemann). The D1/D2 domain of the large-subunit (28S/LSU) rRNA gene was extracted as the first 700 bp (strand-aware) of the longest barrnap-called 28S_rRNA feature per assembly. The internal transcribed spacer region (ITS1–5.8S–ITS2) was extracted with ITSx v1.1.3 (-t Fungi), run on a 1000-bp-padded window around the barrnap-called rRNA locus rather than the whole assembly because the ITSx HMMER pipeline enforces a 100 kb single-sequence input limit incompatible with whole-contig input (Bengtsson-Palme, et al. 2013). Where ITSx returned no confident ITS boundary, the raw interval between the 18S and 28S barrnap coordinates was used as a fallback.

#### Barcode-based species screening

Extracted ITS and D1/D2 LSU marker sequences were queried against the NCBI nucleotide database (nt) with blastn (BLAST+ v2.17.0; (Camacho, et al. 2009)) in remote mode (-outfmt “6 qseqid sseqid pident length mismatch gapopen qstart qend sstart send evalue bitscore qcovs stitle”, -max_target_seqs 5, with a 15 s courtesy delay between queries). For each marker, the highest-bitscore hit whose subject title parsed as a valid binomial (first two tokens: a capitalized alphabetic genus and a lowercase alphabetic specific epithet) was taken as the top hit; non-taxonomic records (*e.g.*, PDB structure entries, “Uncultured…”, “Mutant…” descriptions) were excluded by this filter. This exclusion was consequential: for YH115, a spurious top hit to a PDB structure title (“Chain B2,”) would otherwise have downgraded a correct identification to a false marker disagreement. Percent identity and query coverage were recorded for each marker’s top hit.

#### Whole-genome ANI

Because rDNA barcodes cannot resolve species within the *Saccharomyces sensu stricto* complex (Peter, et al. 2018), whole-genome ANI was used as a confirmatory and, for that clade, authoritative step. Reference genomes were retrieved with the NCBI datasets CLI v18.32.0 (Sayers, et al. 2026), preferring the RefSeq-flagged reference or representative assembly per species (--reference), with fallback to the best available assembly where no RefSeq reference existed. ANI was computed with skani v0.3.2 (dist mode) using a screening threshold of -s 70 (default 80) and the --slow preset (-c 30 sketch compression, *vs.* the default -c 125) (Shaw and Yu 2023). This non-default configuration was necessary because during method development, skani’s default -s 80 threshold was found to silently omit reference comparisons below the k-mer screening cutoff with no error emitted. Several genuine, informative cross-species comparisons in the 79–89% ANI range were being dropped, and lowering the threshold recovered them without altering any species call. For isolates whose barcode hits indicated genus *Saccharomyces*, ANI was computed against a fixed panel of all eight recognized *sensu stricto* species (*S. cerevisiae*, *S. paradoxus*, *S. mikatae*, *S. kudriavzevii*, *S. arboricola*, *S. eubayanus*, *S. uvarum*, and *S. jurei*) rather than only the top barcode hit. A 95% ANI threshold was applied as the species boundary.

#### Consensus species assignment

Species calls combined barcode and ANI evidence. For non-*Saccharomyces* isolates, agreement between the ITS and LSU top hits (each ≥ 90% identity and ≥ 80% query coverage) together with ANI ≥ 95% to the corresponding reference yielded a high-confidence call. Marker disagreement, a single confident marker, or no confident marker produced medium- or low-confidence calls. For *Saccharomyces sensu stricto* isolates, the barcode was treated as uninformative for species-level resolution, and the ANI result was authoritative (Peter, et al. 2018). A single reference clearing the 95% boundary with no other reference within 0.5% of the top ANI value yielded a high-confidence call, whereas multiple references clearing the boundary within 0.5% of one another were flagged as ambiguous or possible hybrids.

#### Mixed-culture resolution (isolate YH140)

YH140 returned discordant ITS and LSU barcode identities (*L. thermotolerans* and *T. delbrueckii*, respectively), an assembly (19.9 Mb) roughly twice the expected single-genome size, and was investigated as a possible mixed culture. Assembly contigs were binned by best match to each candidate reference with minimap2 v2.31-r1302 (-x asm10) (Li 2018), summing per-contig aligned bases (PAF matching-bases column) and assigning each contig to the higher-scoring reference (minimum 10% of contig length aligned; otherwise unassigned). Read-level support was quantified by mapping filtered reads to each reference independently (minimap2 -ax map-ont) and counting primary alignments (samtools v1.23.1, view -F 0×904) (Danecek, et al. 2021). BUSCO (saccharomycetes_odb10) was run independently on each contig bin (Manni, et al. 2021). The approach was designed to distinguish true co-culture (expected to show comparable read-level support for both organisms and to yield two near-complete, non-duplicated single-species bins) from index cross-talk on a multiplexed run, which would produce only a small minority fraction for the second organism. YH140 yielded 53.3% of the assembly (10.6 Mb, 15 contigs) assigned to *L. thermotolerans* and 46.2% (9.2 Mb, 13 contigs) to *T. delbrueckii*, with 78.9% and 41.2% of filtered reads (not mutually exclusive, owing to conserved sequence) mapping to each reference, and each bin reaching > 99.7% BUSCO completeness with < 0.15% duplication, consistent with genuine co-culture.

#### Species re-identification (isolate YH156)

YH156’s standard-pipeline assembly was severely fragmented (426 kb, 54 contigs, 0% BUSCO) and initially resembled a failed run. Both the fragmentation and the 0% score were subsequently attributed to identifiable artifacts rather than to sample or sequencing quality (Part B). Its identity was resolved by the ANI methodology above. An initial candidate identification of *S. japonicus* was tested and rejected: ANI against the *S. japonicus* reference (strain yFS275, GCF_000149845.2) was 87.94% (align-fraction-query 56.02%), well below the 95% boundary. blastn (-remote) of poorly-matching contigs returned consistent high-identity hits (96-99.8% identity, 100% query coverage) to an assembly labeled *S. versatilis* (nom. inval.) strain CBS 103 (GCA_032882995.1; the type strain; a complete 2023 ONT+PacBio+Illumina assembly, 15,482,186 bp across three chromosomes). ANI against this genome was 99.14% (align-fraction-query 71.95%), decisively above the boundary. YH156 is therefore reported as *S. versatilis* (nom. inval.); “nom. inval.” (*nomen invalidum*) indicates that the name has not yet been validly published under the applicable nomenclatural code (Brysch-Herzberg, et al. 2024).

### Part B - Genome assembly, species correction, and structural characterization of *S. versatilis* isolate YH156

#### Diagnosis of the initial assembly failure and assembler comparison

The initial fragmentation was traced to Flye’s (Kolmogorov, et al. 2019) coverage-based contig filter: an rDNA tandem-repeat array at extreme copy number (a single contig at 26,208× coverage, *vs*. 100–800× genome-wide) inflated the genome-wide mean coverage used as the filter’s reference point, causing legitimately assembled single-copy contigs to be discarded as low-coverage outliers. Re-running Flye with --scaffold reproduced the failure, confirming a repeat-graph resolution limit rather than a coverage or filtering issue alone. Two algorithmically distinct assemblers were therefore run in parallel on the pooled reads: Canu v2.3 (development build; git r10515, commit 2a41e93; genomeSize=12m) (Koren, et al. 2017) and hifiasm v0.25.0-r726 (--ont) (Cheng, et al. 2021). Canu produced 211 contigs totaling 17,065,046 bp (N50 163 kb) and hifiasm 214 contigs totaling 20,229,027 bp (N50 280 kb). Canu was selected as the primary draft for its smaller size inflation relative to the reference and the closer agreement between its optimistic and conservative reference-coverage estimates.

#### Chimera detection and correction

Two chimeric contigs were identified independently in both the Canu and hifiasm assemblies, breaking at identical reference coordinates (cross-assembler concordance indicating a real, repeat-driven misassembly rather than assembler-specific noise). Chimeric junctions were verified by assembly-to-reference alignment (minimap2 v2.31-r1302, -x ‘’asm5) (Li 2018), split at the verified breakpoints with seqkit (Shen, et al. 2016), and the corrected pieces reincorporated, yielding a de-chimerized draft of 213 contigs (17,065,046 bp; total length unchanged, confirming no sequence loss).

#### Polishing

The de-chimerized draft was consensus-polished with medaka v2.2.1 (model r1041_e82_400bps_sup_v5.2.0, selected to match the R10.4.1 super-accuracy basecalling), giving the canonical assembly used for all downstream analyses.

#### Completeness assessment and the lineage-database ceiling

Completeness was assessed with BUSCO v5.8.3 (euk_genome mode; metaeuk v7.bba0d80 predictor; hmmsearch v3.4) (Levy Karin, et al. 2020; Manni, et al. 2021) against ascomycota_odb12.2 (2026-05-13 build; 102 genomes; 2,557 BUSCO groups), the most specific valid lineage available for this Taphrinomycotina taxon. (BUSCO’s automatic lineage downloader mis-constructs URLs for versioned lineage names; the dataset was therefore downloaded manually and BUSCO run with --offline.) The polished draft scored 55.4% complete (C:55.4%[S:52.8%,D:2.5%],F:7.5%,M:37.1%). To distinguish assembly incompleteness from a lineage-database limitation, two finished published genomes were scored under identical settings as controls: the *S. japonicus* reference (GCF_000149845.2) scored 57.1%, and the correct-species *S. versatilis* CBS 103 reference (GCA_032882995.1) scored 57.9%. The close clustering of all three values indicates that the ∼42% of BUSCO groups scored missing or fragmented reflects the poor fit of the ascomycota_odb12.2 ortholog set to this early-diverging lineage rather than draft incompleteness. The residual ∼2.5-point gap between the draft and the finished same-species genome represents the draft’s actual incompleteness, consistent with a 213-contig assembly.

#### Species confirmation

The *S. versatilis* identification (Part A) was re-derived independently in this analysis from the polished draft (skani dist -s 70 –slow; (Shaw and Yu 2023)): 87.94% ANI to *S. japonicus* (GCF_000149845.2) *vs.* 99.14% to *S. versatilis* CBS 103 (GCA_032882995.1). Merged, non-redundant minimap2 coverage (-x asm10) (Li 2018) of the draft against the corrected reference was 93.7%, 93.7%, and 94.3% for chromosomes CP130697.1, CP130698.1, and CP130699.1, respectively.

#### Whole-genome structural comparison

For structural comparison to the *S. versatilis* CBS 103 reference, draft contigs were oriented and concatenated into a reference-guided pseudo-chromosome assembly. Structural variants were called with MUMmer4 v4.0.1 (alignment) (Marcais, et al. 2018) followed by SyRI v1.7.1 (Goel, et al. 2019), and visualized with plotsr v1.1.1 (run in a dedicated environment with numpy 1.26.4, pandas 1.5.3, and matplotlib 3.7.3 for tool compatibility) (Goel and Schneeberger 2022). SyRI resolved 99 syntenic regions alongside 13 inversions, 103 translocations, and numerous duplications and smaller variants genome-wide. Three large, well-isolated inversions and one intra-chromosomal translocation were designated high-confidence after visual inspection and a contig-boundary artifact check (below). Two methodological caveats constrain interpretation and are reported with the results: 1) because each contig was reverse-complemented to its dominant reference-alignment strand before concatenation, a real inversion spanning exactly one whole contig would be silently corrected and is a blind spot of this pseudo-chromosome approach; and 2) 20-30% of SyRI’s inversion/translocation/duplication calls fell within ±2 kb of a contig junction (*vs*. a ∼5% random-placement baseline), so a minority of calls are likely placement artifacts, the reason only the largest, cleanest, interior events were treated as high-confidence. Confident chromosome-scale confirmation of these events would require higher-contiguity data (*e.g.*, Hi-C or ultra-long reads).

#### Repeat and transposable-element characterization

A complex duplication/translocation signal on chromosome 2 was characterized as dispersed repeat content. Contigs flagged by Canu’s internal sugRept heuristic and showing coverage 4–6× the genome-wide median (8.96×) were extracted (Koren, et al. 2017). Nucleotide BLAST (blastn vs. nt) of representative windows returned near-identical (98-99.9%) matches to multiple different chromosomes of the *S. versatilis* reference, and translated BLAST (blastx *vs*. nr, restricted to Fungi) identified homology to Tf1/Tf2-type retrotransposon polyproteins (*e.g.*, *S. pombe* “Transposon Tf1-107 polyprotein,” 39.0% identity, e = 2.2 × 10⁻¹DD). The signal was therefore attributed to a dispersed Tf1/Tf2-type LTR retrotransposon family conserved across *Schizosaccharomyces* (Levin 1995), rather than to an assembly artifact.

The YH156 assembly is reported throughout as a draft (213 contigs, unscaffolded, with residual repeat-region ambiguity), not a finished reference-quality genome.

### Part C - Population-genomic placement and diversity analysis

The four isolates confirmed as *S. cerevisiae* (YH123, YH166, YH196, and YH229) and the wild isolate DoF1 (Gray, et al. 2026) were placed within a reference population panel and characterized for heterozygosity, copy-number variation, and interspecific hybrid ancestry.

#### Reference population panel

Isolates were placed against the *S. cerevisiae* population-genomics resource of (Loegler, et al. 2024), comprising 3,034 classified isolates (3,039 total sequenced samples) organized into 39 clades within four superclades (Wine, Beer, Asian Fermentation, Wild) plus 321 unassigned/admixed isolates. The panel’s S288C R64 reference FASTA (chromosomes chromosome1–chromosome16) was used for all alignments to maintain a shared coordinate system, together with the panel’s joint-genotyped, quality-filtered SNP catalog (1,916,611 biallelic SNP sites; filters: depth ≥ 10, genotype quality ≥ 20, ≥ 99% non-missing, excess-heterozygosity threshold), both obtained from the panel’s Zenodo repositories (10.5281/zenodo.12580561, 10.5281/zenodo.12571280). Per-isolate metadata (clade/superclade, ploidy, zygosity, heterozygosity, geographic and ecological origin) were drawn from the panel’s Supplementary Table 1.

#### Read alignment and variant calling

Reads for each of the five isolates were aligned to the panel’s S288C R64 reference with minimap2 v2.31-r1302 (map-ont) (Li 2018), with read-group sample tags assigned during alignment. Variants were called per isolate with Clair3 v2.0.2 (Zheng, et al. 2022) using the r1041_e82_400bps_sup_v500 pretrained model (the closest available match to the data’s v4.3.0 sup basecall tag). The --include_all_ctgs flag was required because Clair3’s default contig-name filtering, tuned for human chromosome naming, otherwise silently excludes non-standard contig names such as “chromosome1,” producing empty callsets despite successful alignment. This is noted as a reproducibility pitfall.

#### Site-subsampled phylogenetic placement

To place the five isolates within the panel tractably, 50,437 evenly spaced sites (every 38th site) were subsampled from the 1,916,611-site catalog. Panel genotypes at these sites were extracted from the joint VCF, and the study isolates were genotyped at the same positions by combining: 1) their Clair3 (Zheng, et al. 2022) calls where present and 2) samtools v1.23.1 (Danecek, et al. 2021) depth-based confirmation (≥ 5× read depth) of homozygous-reference status at non-variant panel sites, an approach adopted because classical genotype-likelihood callers (bcftools v1.23.1 mpileup/call) proved computationally prohibitive at this scale on a 12-Mb genome at high per-sample depth. The combined 3,039 × 50,437 genotype matrix was converted to a multi-sample FASTA alignment, encoding heterozygous genotypes as IUPAC ambiguity codes and excluding indel and multiallelic sites, yielding 46,705 usable biallelic SNP sites. An approximately-maximum-likelihood tree was inferred with FastTree v2.2.0 (double precision; nucleotide mode; Jukes–Cantor model) (Price, et al. 2010); reported node supports are FastTree’s local Shimodaira–Hasegawa-like values, not bootstraps. Clade and superclade placement was determined by examining each isolate’s nearest neighbors at increasing tree radii (Biopython v1.87 Phylo; (Cock, et al. 2009)) and cross-referencing neighbor identities against the panel’s clade metadata.

#### Heterozygosity and private-variant analysis

Per-isolate heterozygosity was quantified as the count of heterozygous SNP genotypes (full, non-subsampled Clair3 callset; (Zheng, et al. 2022)) divided by the reference genome length (12,071,326 bp) and compared against panel per-superclade mean heterozygosity (Supplementary Table 1). Private variation was quantified as the fraction of each isolate’s called SNPs at positions absent from the 1,916,611-site catalog, with the explicit caveat that this measure conflates genuine novel variation with sites excluded by the panel’s cohort-level quality filters and platform-specific calling differences between the panel’s Illumina/GATK pipeline and this study’s Nanopore/Clair3 pipeline. It is therefore not interpreted as a clean novelty estimate.

#### Superclade allele-sharing analysis (isolate YH166)

For YH166, whose *S. cerevisiae* subgenome did not resolve into any panel clade, a superclade-level allele-sharing analysis was performed. At YH166’s heterozygous sites also present in the panel catalog (n = 35,800), the mean alternate-allele frequency within each superclade was computed to test whether one superclade’s alleles were disproportionately represented among heterozygous sites.

#### Copy-number and aneuploidy screening

Chromosome-level relative copy number was assessed with samtools coverage (Danecek, et al. 2021), normalizing each chromosome’s mean depth to the isolate’s genome-wide mean. Finer-scale variation was assessed with mosdepth v0.3.14 (Pedersen and Quinlan 2018) in 1 kb windows; outlier windows (> 2.0× or < 0.3× the genome-wide mean, in runs of ≥ 3 consecutive windows) were flagged and cross-referenced across the five isolates. Outlier regions recurring at matching coordinates in ≥ 3 of 5 isolates (114 regions) were classified as shared reference-mapping artifacts (repetitive and subtelomeric regions) rather than biological variation.

#### Screening for interspecific *S. eubayanus* ancestry (isolates YH166 and YH229)

Two isolates, YH166 and YH229, showed reduced primary read-mapping rates to the S288C reference (60.5% and 77.6%, respectively, *vs*. 93–96% for the other three *S. cerevisiae* isolates), prompting a test for interspecific *S. eubayanus* ancestry. Reads were competitively mapped (minimap2 v2.31-r1302, map-ont, --secondary=no; (Li 2018)) against a cytoplasm-aware combined reference comprising the *S. cerevisiae* S288C 16 nuclear chromosomes plus the *S. cerevisiae* mitochondrion (NC_001224.1) and 2-micron plasmid (NC_001398.1), together with the *S. eubayanus* FM1318 16 nuclear chromosomes (its own organellar and unplaced-scaffold contigs excluded because dedicated two-contig comparisons found no evidence of retained *S. eubayanus* cytoplasmic material). Including the *S. cerevisiae* cytoplasmic elements is essential rather than optional. The standard S288C nuclear assembly omits both (Engel, et al. 2014), so under a nuclear-only reference, the isolates’ own organellar and plasmid reads have no legitimate target, mismap onto nuclear sequence, and inflate apparent interspecific ancestry. Depth was summarized in 10-kb windows (mosdepth v0.3.14; bedtools v2.31.1 for window generation and block merging) (Pedersen and Quinlan 2018; Quinlan and Hall 2010). A window was scored *S. eubayanus*-derived at ≥5× mean depth, the genome-wide fraction was reported as candidate *S. eubayanus* bp ÷ total callable nuclear bp at ≥5× across both species, and relative dosage as the median depth of candidate windows divided by the genome-wide median depth of *S. cerevisiae*-side windows.

Candidate signal was then tested against matched negative controls. Two wild *S. cerevisiae* with no plausible *S. eubayanus* contact – YH123 and DoF1, sequenced on the same platform, chemistry, and basecaller – were processed through the identical pipeline, and isolates and controls were down-sampled (samtools view -s; five random seeds per condition; (Danecek, et al. 2021)) to common depths. Four criteria were applied to every candidate block: 1) survival of a terminus mask excluding the first and last 20 kb of each nuclear chromosome; 2) absence of the same coordinates from the control set at matched depth; 3) a consistent *S. eubayanus* preference on per-read realignment against single-species references; and 4) representation in the isolate’s *de novo* assembly (minimap2, -x asm10; (Li 2018)). No block in either isolate satisfied all four criteria. Full parameters, control results, and the derivation of the false-positive floor are reported in the companion methods study (companion paper, Taylor *et al*.).

#### Visualization

A circular cladogram of all 3,039 samples was rendered in Python (matplotlib v3.11.0; (Hunter 2007)) using an equal-angle radial layout with node depth (topological, unit branch length) determining radius. Branches were colored by the dominant superclade of their descendant tip set, and superclade regions were shaded using the angular bounds of each superclade’s largest contiguous tip block(s), with a minimum contiguous-block size of 6 tips for named superclades and 10 tips for the unassigned/admixed group (an empirical decluttering threshold, not a statistical criterion).

### Part D - *L. thermotolerans* placement

Three *L. thermotolerans* isolates (YH26, YH72, and the *L. thermotolerans* component of the YH140 co-culture) were placed relative to the 145-strain population panel of (Vicente, et al. 2025), which resolves six ecological/geographic clusters (Asia, Americas, Canada-trees, Europe/Domestic-1, Europe/Domestic-2, Europe-Mix). Because the panel’s filtered genotype matrix could not be obtained from the source authors within the analysis window, the reference panel was reconstructed independently from the published raw reads (NCBI BioProject PRJNA1111406; ENA PRJEB29656): reads were aligned to the *L. thermotolerans* CBS 6340 reference with bwa-mem2 v2.3 (Vasimuddin, et al. 2019), and variants were jointly called across the 145 panel strains with bcftools v1.23.1 (mpileup/call, per-chromosome), normalized (norm - m -any), and hard-filtered to biallelic SNPs (QUAL≥30 && MQ≥40 && |MQBZ|≤12.5 && |RPBZ|≤8) (Danecek, et al. 2021), retaining 93.0% of records, comparable to the 93.9% retention reported by the source study. This reconstruction used bcftools-native annotations (MQBZ, RPBZ) as analogs of the source study’s GATK-based statistics (MQRankSum, ReadPosRankSum), and is therefore methodologically comparable to, but not a byte-for-byte reproduction of, the published calls. The source study’s authoritative filtered VCF was subsequently obtained, and the ADMIXTURE (Alexander, et al. 2009) and PCA placement analyses were independently re-run against it. Both were concordant with the reconstruction (99.995% Canada-trees ancestry in both; PCA placement nearest the Canada-trees centroid in both, with nearly identical variance explained).

For the three study isolates, reads were aligned to CBS 6340 with bwa-mem2 (Vasimuddin, et al. 2019); for the YH140 co-culture, reads were assigned to *L. thermotolerans* by retaining primary alignments at MAPQ ≥ 20 against a *Lachancea*-only reference (57,944 reads; the additional MAPQ filter accounts for the modest reduction relative to the primary-only co-culture count in Part A). The study isolates were merged with the reconstructed panel (148 samples) and placed by three independent methods. Supervised ADMIXTURE v1.3.0 was run at K = 6 with the panel clusters as reference populations (Alexander, et al. 2009). Principal-component analysis was performed with SNPRelate v1.36.0 (gdsfmt v1.38.0) (Zheng, et al. 2012) in R v4.3.3 on a PLINK-filtered SNP set (--geno 0.1 --maf 0.01; PLINK v1.9.0-b.8) (Chang, et al. 2015). A BIONJ tree was inferred from a genome-wide distance matrix with ape v5.8-1 (Paradis and Schliep 2019); monophyly of each panel cluster was tested with ape::is.monophyletic.

Figures were rendered with ggplot2 v3.5.2 (Villanueva and Chen 2019) and ggtree v3.10.0 (Yu, et al. 2017). Population-genomic steps (ADMIXTURE, PLINK, R and its packages) were run in a dedicated popgen conda environment; read alignment and variant calling used the project yeast-id environment.

## Software and data availability

Analyses used seqkit v2.13.0, NanoPlot v1.47.1, chopper v0.13.0, Flye v2.9.6-b1802, Canu v2.3 (git r10515, commit 2a41e93), hifiasm v0.25.0-r726, medaka v2.2.1, BUSCO v5.8.3 (miniprot v0.18-r281, metaeuk v7.bba0d80, hmmsearch v3.4), barrnap v1.10.6, ITSx v1.1.3, blastn/blastx (BLAST+ v2.17.0), NCBI datasets CLI v18.32.0, skani v0.3.2, minimap2 v2.31-r1302, samtools/bcftools/htslib v1.23.1, Clair3 v2.0.2, FastTree v2.2.0, mosdepth v0.3.14, MUMmer4 v4.0.1, SyRI v1.7.1, plotsr v1.1.1, Biopython v1.87, and matplotlib (v3.11.0; v3.7.3 in the synteny environment) unless otherwise noted. Population-genomic placement (Part D) additionally used bwa-mem2 v2.3, ADMIXTURE v1.3.0, PLINK v1.9.0-b.8, and R v4.3.3 with SNPRelate v1.36.0, gdsfmt v1.38.0, ape v5.8-1, ggtree v3.10.0, and ggplot2 v3.5.2 unless otherwise noted. Analyses were distributed across three conda environments: yeast-id (assembly, alignment, variant calling, ANI, phylogenetics), synteny (whole-genome structural comparison), and popgen (ADMIXTURE, PLINK, R and its packages). Reference genomes were obtained from NCBI (accessions cited in text and Table S1). The *S. cerevisiae* population panel and its variant catalog were obtained from Zenodo (10.5281/zenodo.12580561, 10.5281/zenodo.12571280); the *L. thermotolerans* panel was reconstructed from published raw reads under NCBI BioProject PRJNA1111406 and ENA PRJEB29656 (see Part D). DoF1 reads are available under NCBI BioProject PRJNA1477971 (run SRR39135318).

## DATA AVAILABILITY

Raw sequencing reads and genome assemblies generated in this study have been deposited in the NCBI Sequence Read Archive (SRA) and GenBank under BioProject accession PRJNA[still processing] (SRA accession SUB16383871). The previously published DoF1 sequencing data are available under BioProject PRJNA1477971 (SRA accession SRR39135318). Reference genome and isolate accessions are listed in Tables 1 and S1. Analysis scripts and the complete bioinformatic workflow for this study are available at Zenodo (DOI: 10.5281/zenodo.21811551); the introgression-withdrawal analysis (YH166, YH229) used the FIDDL toolkit, available separately at 10.5281/zenodo.21509543. All additional data supporting the findings of this study, including phylogenetic trees, variant datasets, and supplementary tables, are provided within the article, its Supplementary Material, or the associated public repositories.

## ACKNOWLEDGMENTS

We thank Kris Brown, Kara Osburn, Justin Miller, and David Nickens for participating in our yeast hunting efforts. This work was supported by funds from Plasmidsaurus and the Walter Center for Career Achievement. This work is also dedicated to the memory of David T. Bochman (1949-2025), on whose land YH72 was isolated.

## AUTHOR CONTRIBUTIONS

**Kennadi A. Shumaker:** Conceptualization, Formal analysis, Investigation, Methodology, Resources, Validation, Visualization, Writing – review and editing. **Kaitlyn Taylor:** Conceptualization, Formal analysis, Investigation, Methodology, Resources, Validation, Visualization, Writing – review and editing. **Spencer J. Gray:** Conceptualization, Formal analysis, Investigation, Methodology, Resources, Validation, Visualization, Writing – review and editing. **Matthew L. Bochman:** Conceptualization, Data curation, Formal analysis, Investigation, Methodology, Resources, Software, Project administration, Resources, Supervision, Validation, Visualization, Writing – original draft, Writing – review and editing.

