## Supplementary materials for "Genome-scale characterization of wild yeasts reveals cryptic diversity and population structure across three genera"

**Supplementary Tables**

**Table S1. Reference genomes and population panels used for identification and placement.**

**Supplementary Figures**

**Figure S1. Contig binning of the YH140 co-culture.**

**Figure S2. Per-chromosome reference coverage (%).**

**Figure S3. A BUSCO completeness ceiling for deep-branching fission yeasts.**

**Figure S4. Structural-variant summary for YH156 *vs*. CBS 103.**

**Supplementary References**

**Table S1. Reference genomes and population panels used for identification and placement.** Every reference assembly used in the ANI panels, structural comparison, introgression mapping, and phylogenomic placement, with species, strain, assembly accession, source database, and the analysis in which it was used. Pinned references include *S. cerevisiae* S288C (GCF_000146045.2) with its mitochondrial genome (NC_001224.1) and 2-micron plasmid (NC_001398.1); *S. japonicus* (GCF_000149845.2); *S. versatilis* CBS 103 (GCA_032882995.1); *S. eubayanus* FM1318 [confirm assembly accession]; the two *S. pastorianus* references (CBS 1483, GCA_011022315.1; CBS 1513, GCA_056824455.1); and *L. thermotolerans* CBS 6340 [confirm accession]. Population panels: the 3,034-genome *S. cerevisiae* panel and variant catalog (Loegler et al., 2024) (Zenodo 10.5281/zenodo.12580561, 10.5281/zenodo.12571280) and the 145-strain *L. thermotolerans* panel (Vicente et al., 2025) (NCBI BioProject PRJNA1111406; ENA PRJEB29656), the latter reconstructed from raw reads (Methods, Part D).

| **#** | **Species** | **Strain** | **Assembly accession** | **Source** | **Used in** |
| --- | --- | --- | --- | --- | --- |
| 1 | *Saccharomyces cerevisiae* | S288C | GCF_000146045.2 | RefSeq | *sensu stricto* ANI panel (Part A); hybrid-detection *S. cerevisiae* reference (Part C) ᵃ |
| 2 | *Saccharomyces paradoxus* | CBS432 | GCA_002079055.1 | GenBank | *sensu stricto* ANI panel; hybrid-detection |
| 3 | *Saccharomyces mikatae* | IFO1815 | GCF_947241705.1 | RefSeq | *sensu stricto* ANI panel; hybrid-detection |
| 4 | *Saccharomyces kudriavzevii* | CR85 | GCA_900682695.1 | GenBank | *sensu stricto* ANI panel; hybrid-detection |
| 5 | *Saccharomyces arboricola* | H-6 | GCF_000292725.1 | RefSeq | *sensu stricto* ANI panel; hybrid-detection |
| 6 | *Saccharomyces uvarum* | CBS7001 | GCA_027557585.1 | GenBank | *sensu stricto* ANI panel; hybrid-detection |
| 7 | *Saccharomyces jurei* | C1003 (= NCYC 3947) | GCA_900290405.1 [ | GenBank | *sensu stricto* ANI panel; hybrid-detection |
| 8 | *Lachancea thermotolerans* | CBS 6340 | GCF_000142805.1 | RefSeq | species-ID ANI (YH26, YH72); YH140 contig binning; hybrid-detection outgroup |
| 9 | *Torulaspora delbrueckii* | CBS 1146 | GCF_000243375.1 | RefSeq | YH140 contig binning; hybrid-detection outgroup |
| 10 | *Schizosaccharomyces pombe* | 972h | GCF_000002945.1 | RefSeq | *Schizosaccharomyces* ANI panel (YH115); hybrid-detection outgroup |
| 11 | *Schizosaccharomyces japonicus* | yFS275 | GCF_000149845.2 | RefSeq | *Schizosaccharomyces* ANI panel; YH156 comparison/control |
| 12 | *Schizosaccharomyces octosporus* | yFS286 | GCF_000150505.1 | RefSeq | *Schizosaccharomyces* ANI panel (YH156 re-ID) |
| 13 | *Schizosaccharomyces cryophilus* | OY26 | GCA_000004155.2 | GenBank | *Schizosaccharomyces* ANI panel (YH156 re-ID) |
| 14 | *Schizosaccharomyces versatilis* (nom. inval.) | CBS 103 | GCA_032882995.1 | GenBank | YH156 species assignment; whole-genome structural comparison |
| 15 | *Saccharomyces osmophilus* | CBS 15793 | GCA_027921745.1 | GenBank | hybrid-detection outgroup |
| 16 | *Saccharomyces eubayanus* | FM1318 | GCF_001298625.1 | RefSeq | *sensu stricto* ANI panel; hybrid-detection (subgenome of YH166, YH229); *S. eubayanus* introgression quantification and subgenome placement (YH166, YH229) ᶜ |
| 17 | *Saccharomyces cerevisiae* (mitochondrion) | S288C | NC_001224.1 | RefSeq | mitochondrial-inheritance test (extracted from GCF_000146045.2) |
| 18 | *Saccharomyces cerevisiae* (2-micron plasmid) | — | NC_001398.1 | RefSeq | 2-micron plasmid-inheritance test |
| 19 | *Saccharomyces pastorianus* | CBS 1483 (Group 2/Frohberg) | GCA_011022315.1 | GenBank | lager *Sc*/*Se* chimeric-junction architecture comparison |
| 20 | *Saccharomyces pastorianus* | CBS 1513 (Group 1/Saaz) | GCA_056824455.1 | GenBank | lager *Sc*/*Se* chimeric-junction architecture comparison |

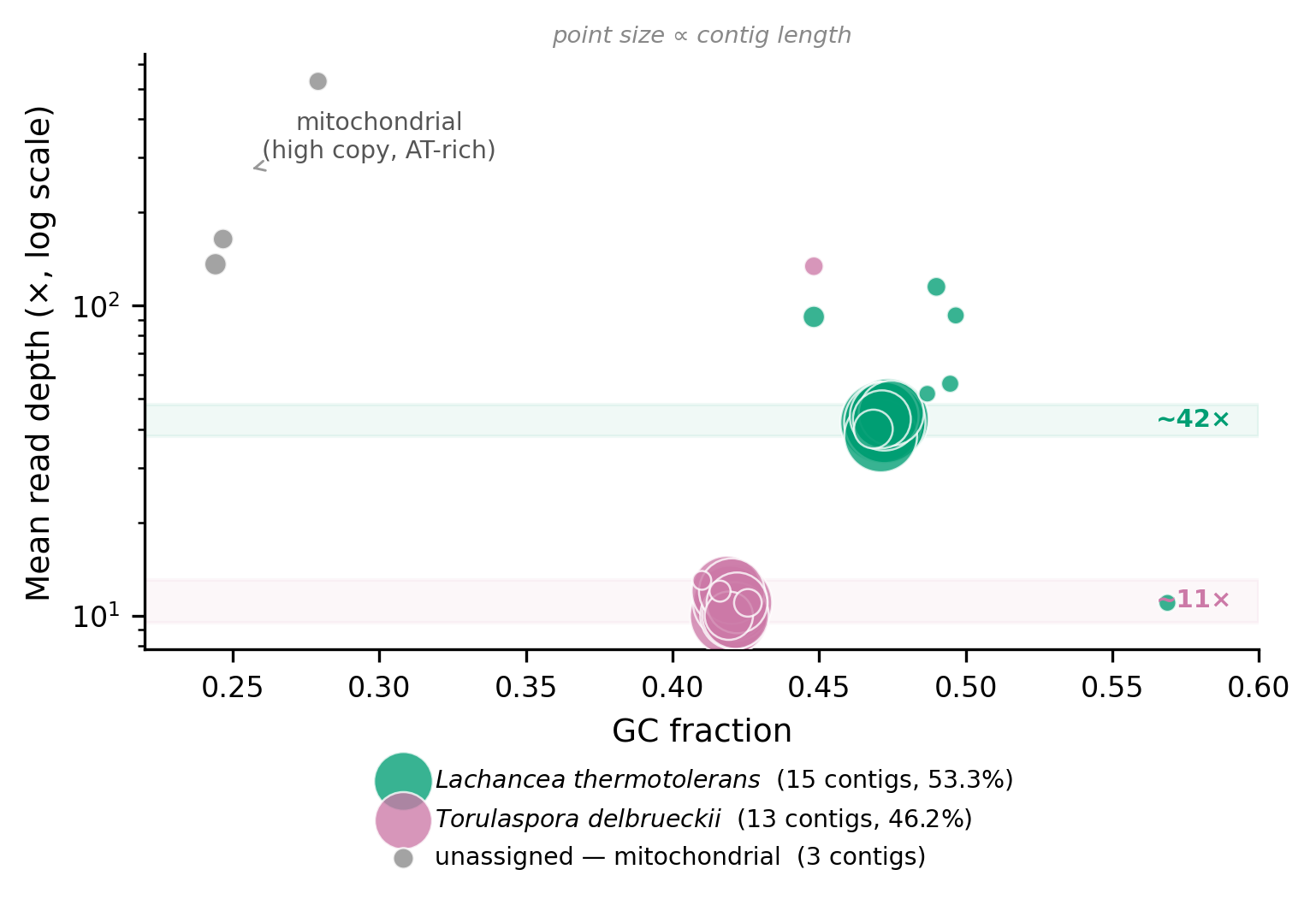

**Figure S1. Contig binning of the YH140 co-culture.** Per-contig mean read depth *vs*. GC fraction for the YH140 assembly (point area proportional to contig length). Contigs were assigned to *Lachancea thermotolerans* (15 contigs, 53.3% of the assembly) or *Torulaspora delbrueckii* (13 contigs, 46.2%) by competitive alignment to each reference genome (minimap2 -x asm10; a contig assigned to whichever reference received more aligned bases, ≥10% of contig length required). The two components separate cleanly on depth – *L. thermotolerans* at ~42× and *T. delbrueckii* at ~11×, reflecting their different abundance in the co-culture – with GC providing a secondary, weaker separation. Because assignment was made by alignment rather than by depth or GC, the depth/GC separation is an independent corroboration of the two-species partition rather than a restatement of the binning criterion. Three small, high-depth, AT-rich contigs (GC ~0.25, depth 136–529×) are mitochondrial. Together these confirm a genuine two-species co-culture rather than a single isolate.

|  | Flye | Canu | hifiasm |
| --- | --- | --- | --- |
| Chr 1 | 36.7 | 91.5 | 94.3 |
| Chr 2 | 1.2 | 91.8 | 95.1 |
| Chr 3 | 33.4 | 89.6 | 93.1 |

**Figure S2. Per-chromosome reference coverage (%).** Flye could not resolve chromosome 2 under any configuration tested. Canu was selected for subsequent analyses because it yielded the highest contiguity and lowest duplication.

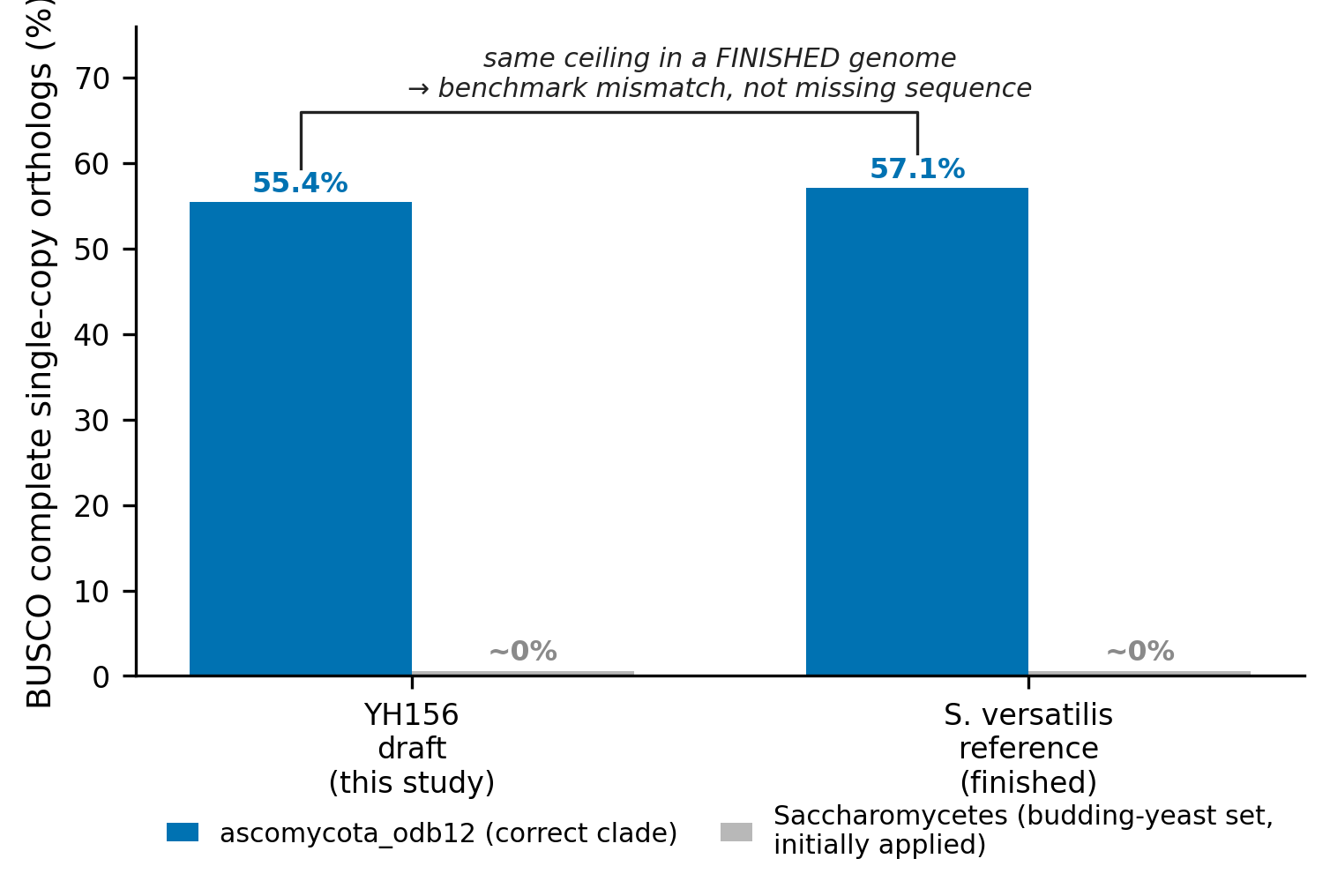

**Figure S3. A BUSCO completeness ceiling for deep-branching fission yeasts.** Single-copy ortholog completeness (ascomycota_odb12.2, BUSCO 5.8.3, --metaeuk) scored identically for the YH156 draft assembly (55.4%) and the finished, conspecific *S. versatilis* CBS 103 reference (57.9%; GCA_032882995.1), contrasted with the ~0% recovered under the budding-yeast (Saccharomycetes) set initially applied. *S. japonicus* (GCF_000149845.2), a congener, scores 57.1% on the same benchmark. All genomes were scored under identical settings, so the comparison is internally consistent. The shared low ceiling in a finished conspecific genome shows that the orthologs scored "missing" in YH156 reflect the benchmark's mismatch to these early-diverging taxa rather than assembly incompleteness.

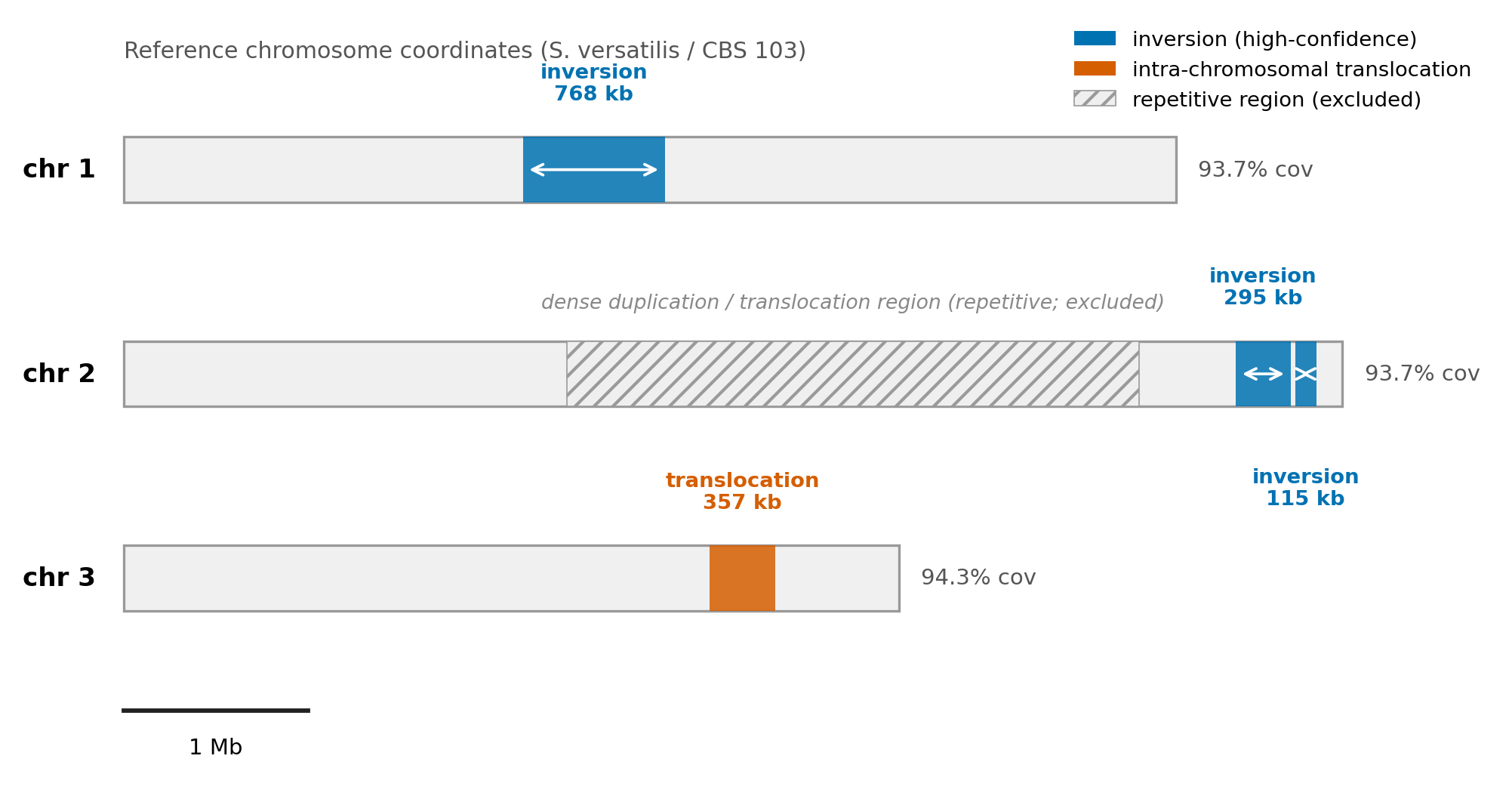

**Figure S4. Structural-variant summary for YH156 *vs*. CBS 103.** The three high-confidence rearrangements underlying Figure 3 — the ~768 kb chromosome-1 inversion (ref ~2,162,341–2,930,379), the two chromosome-2 inversions (~295 kb, ref ~6,024,352–6,319,603; ~115 kb, ref ~6,343,879–6,458,904), and the ~357 kb intra-chromosomal chromosome-3 translocation (ref ~3,174,261–3,530,982) — drawn to scale on their reference chromosomes, with per-chromosome reference coverage of the polished draft (93.7%, 93.7%, 94.3%). Coordinates are per-chromosome; chromosome 2 of this assembly is 7.34 Mb (CP130698.1), roughly double its *S. japonicus* counterpart, so the multi-megabase chromosome-2 coordinates are on-scale rather than concatenated. The dense chromosome-2 duplication/translocation region (~2.4–5.5 Mb), attributed to repetitive sequence, is shown hatched and excluded from the high-confidence set. The draft, contig-level assembly makes individual calls provisional.

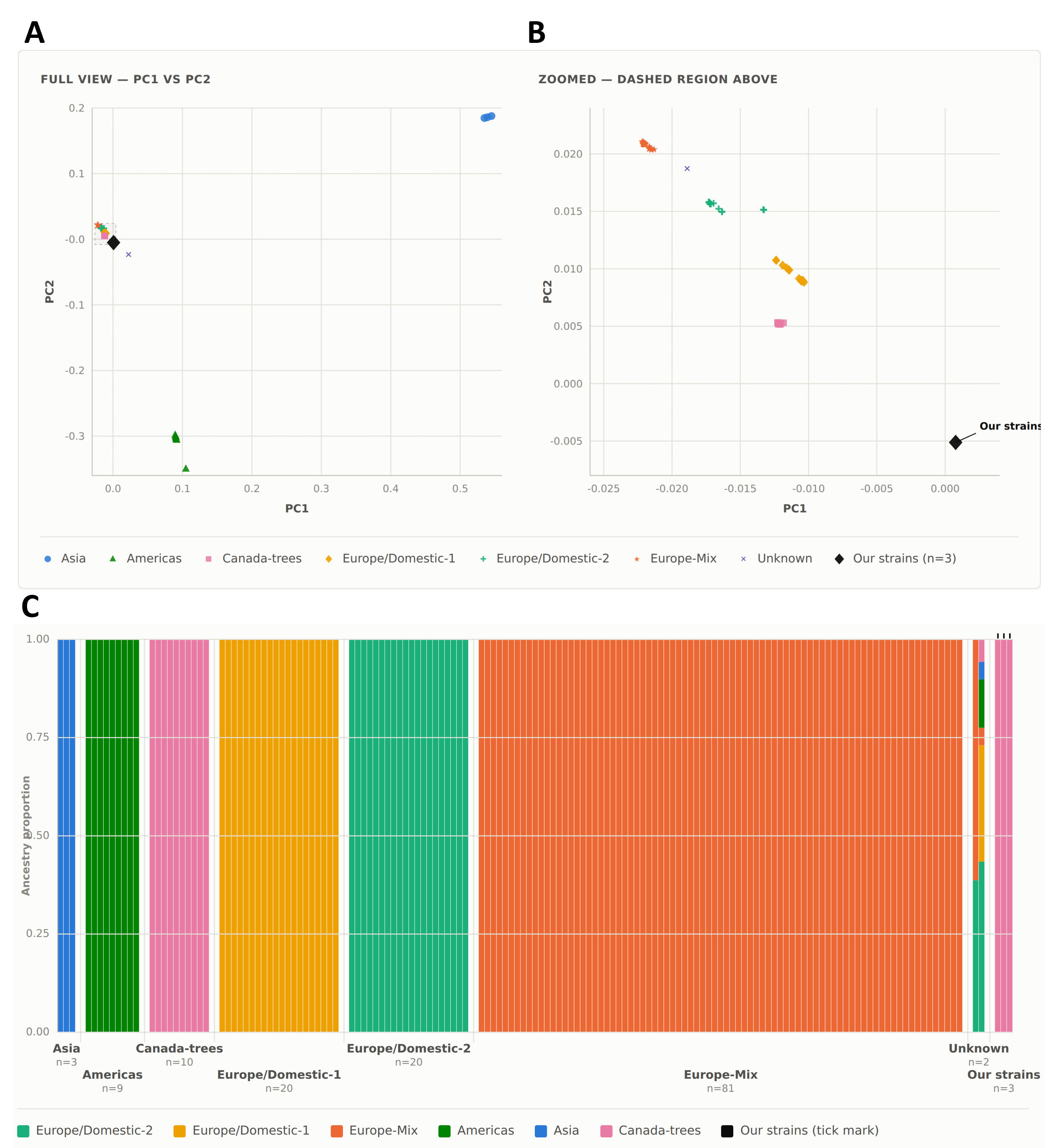

**FigureS5. Placement of *L. thermotolerans* isolates within a 145-strain population panel.** The three collection isolates (YH26, YH72, and the YH140 *L. thermotolerans* bin) placed against the six-cluster panel of (Vicente et al., 2025). (**A**) Principal-component analysis (PC1, 21.80%; PC2, 19.40% of variance) of the 145 panel strains plus the three study strains, colored by ecological/geographic cluster (Asia, n = 3; Americas, n = 9; Canada-trees, n = 10; Europe/Domestic-1, n = 20; Europe/Domestic-2, n = 20; Europe-Mix, n = 81; and Unknown, n = 2); study strains shown as black diamonds (n = 3). (**B**) Zoomed inset of the boxed region in (A): the three study strains form a tight cluster nearest the Canada-trees strains, offset along PC2. (**C**) Supervised ADMIXTURE ancestry proportions (K = 6, panel clusters as reference populations). Nearly every published strain resolves to >99.9% membership in its own labeled cluster. The study strains show 99.995% Canada-trees ancestry. Each bar is one sample’s estimated ancestry proportions across the six published ecological/geographic clusters, with cluster identity fixed from Table S1 as ADMIXTURE’s supervised reference labels as a model-based placement, independent of the PCA and BIONJ-tree cross-checks. Samples are grouped by assigned cluster. The two visibly mixed bars in the Unknown category are the two Table S1 gap samples (no published cluster alignment). One splits Europe/Domestic-2 *vs*. Europe-mix ancestry; the other is admixed across most clusters. The panel was reconstructed from raw reads (Methods, Part D); placement is concordant across PCA, ADMIXTURE, and a BIONJ tree (Figure 4).
